# Large-scale quantification of human osteocyte lacunar morphological biomarkers as assessed by ultra-high-resolution desktop micro-computed tomography

**DOI:** 10.1101/2021.01.05.425223

**Authors:** Elliott Goff, Federica Buccino, Chiara Bregoli, Jonathan P. McKinley, Basil Aeppli, Robert R. Recker, Elizabeth Shane, Adi Cohen, Gisela Kuhn, Ralph Müller

## Abstract

Ultra-high-resolution imaging of the osteocyte lacuno-canalicular network (LCN) three-dimensionally (3D) in a high-throughput fashion has greatly improved the morphological knowledge about the constituent structures – positioning them as potential biomarkers. Technologies such as serial focused ion beam/scanning electron microscopy (FIB/SEM) and confocal scanning laser microscopy (CLSM) can image in extremely high resolution, yet only capture a small number of lacunae. Synchrotron radiation computed tomography (SR-CT) can image with both high resolution and high throughput but has a limited availability. Desktop micro-computed tomography (micro-CT) provides an attractive balance: high-throughput imaging on the micron level without the restrictions of SR-CT availability. Over the past decade, desktop micro-CT has been used to image osteocyte lacunae in a variety of animals, yet few studies have employed it to image human lacunae using clinical biopsies.

In this study, accuracy, reproducibility, and sensitivity of large-scale quantification of human osteocyte lacunar morphometries were assessed by ultra-high-resolution desktop micro-computed tomography. For this purpose, thirty-one transiliac human bone biopsies containing trabecular and cortical regions were imaged using ultra-high-resolution desktop micro-CT at a nominal isotropic voxel resolution of 1.2μm. The resulting 3D images were segmented, component labeled, and the following morphometric parameters of 7.71 million lacunae were measured: Lacunar number (Lc.N), density (Lc.N/BV), porosity (Lc.TV/BV), volume (Lc.V), surface area (Lc.S), surface area to volume ratio (Lc.S/Lc.V), stretch (Lc.St), oblateness (Lc.Ob), sphericity (Lc.Sr), equancy (Lc.Eq), and angle (Lc.θ).

Accuracy was quantified by comparing automated lacunar identification to manual identification. Mean true positive rate (TPR), false positive rate (FPR), and false negative rate (FNR) were 89.0%, 3.4%, and 11.0%, respectively. Regarding the reproducibility of lacunar morphometry from repeated measurements, precision errors were low (0.2 – 3.0%) and intraclass correlation coefficients were high (0.960 – 0.999). Significant differences between cortical and trabecular regions (p<0.001) existed for Lc.N/BV, Lc.TV/BV, local lacunar surface area (<Lc.S>), and local lacunar volume (<Lc.V>), all of which demonstrate the sensitivity of the method and are possible biomarker candidates. This study provides the foundation required for future large-scale morphometric studies using ultra-high-resolution desktop micro-CT and high-throughput analysis of millions of osteocyte lacunae in human bone samples. Furthermore, the validation of this technology for imaging of human lacunar properties establishes the quality and reliability required for the accurate, precise, and sensitive assessment of osteocyte morphometry in clinical bone biopsies.

## 1. INTRODUCTION

Bone as an organ provides humans with the necessary structural support to sustain locomotion and dynamic movement in daily life. The organ is uniquely capable of adapting its structure to meet the mechanical demands that are placed upon it [1]. This adaptation of bone has been described by Roux as bone (re)modeling [2]. Central to this process are the osteocytes: the most abundant bone cell type, embedded deeply within the bone matrix, and each ensconced within individual compartments called lacunae [2, 3]. Woven together by a large number of dendrites that extend from each cell, the lacuno-canalicular network (LCN) is one of the most intricately connected networks in the human body, and the scale is comparable with the network of neurons in the human brain [4]. Compelling studies over the last thirty years have revealed and emphasized the functional importance of the cells and processes within the LCN to sense mechanical signals, to transduce them into chemical signals, and to orchestrate the bone (re)modeling process through guided bone formation and bone resorption [5–10].

After cell death, the fossilized lacuna remains intact, allowing the lacuna’s three-dimensional (3D) geometry to be extracted via several imaging techniques at the sub-micrometer resolution. Today, serial focused ion beam/scanning electron microscopy (FIB/SEM) possesses the highest spatial resolution in the nanometer range. Yet, while this technology allows for features like individual dendritic processes to be resolved, the depth range is a major limitation, and only a few dozen lacunae in the tissue can be captured simultaneously [11–14]. Other researchers have implemented confocal laser scanning microscopy (CLSM) to investigate lacunar geometry in mice [15, 16] and in clinical bone biopsies [17], but again the shallow tissue depth that can be explored is a limitation and hence only small subsections of bone consisting of a few dozen to a few hundred lacunae can be measured with CLSM. Both FIB/SEM and CLSM suffer from a lack of scalability since the time required for a study with more than a few hundred lacunae makes the technologies impractical for any type of large-scale lacunar analysis. Alternatively, several groups have used high-resolution x-ray-based approaches such as synchrotron radiation computed tomography (SR-CT). Several studies, Mader et al. [18] in particular, have been successful in separating the porous lacunae from the surrounding matrix in a high-throughput fashion in complete intact mouse femurs [18–24]. However, SR-CT is an imaging tool that requires access to a beamline facility, of which only a few in the world exist, and hence the availability is limited for most researchers due to timing restrictions. A fourth imaging tool, conventional x-ray based ultra-high-resolution desktop micro-computed tomography (micro-CT), provides a reasonable balance between CLSM and SR-CT. Ultra-high-resolution desktop micro-CT allows for the extraction of millions of lacunae from complete bone biopsies without the need to request approval for limited time slots or experienced personnel at beamline facilities. Furthermore, desktop micro-CT is an established and validated technology that has been implemented for laboratory-based bone research for several decades [25–28]. Therefore, it is necessary that a technique be developed for large-scale, high-throughput imaging of osteocyte lacunar networks in clinical bone biopsies.

Equally as important as the 3D images acquired are the individual structures that are extracted from these images as well as the accurate, reproducible, and sensitive quantification of their morphometry. Specifically, with ultra-high-resolution osteocyte imaging, it is imperative that the lacunar morphometric parameters are well defined and measured accordingly. Great strides have been made towards the standardization of these metrics in recent years, and several studies have explored different basic measures such as lacunar density, shape, and orientation [18, 20, 21, 29–34]. Mader et al. have most thoroughly described and validated both simple and abstract lacunar morphometric parameters, and hence this study follows their naming convention and mathematical definitions [18]. The combination of well-defined lacunar morphometric parameters and a validated imaging and analysis methodology allow for the emergence of biomarkers. These morphometric biomarkers have the potential to be used to differentiate between diseased and healthy bone, old and young bone, or even the region of bone within the body.

This study aims to provide researchers with a reliable method of large-scale lacunar imaging, accurate automated identification, and measurements of each resulting 3D lacunar structure using a technology that is widely accessible – ultra-high-resolution desktop micro-CT. Furthermore, we demonstrate the power of such a high-throughput analysis by measuring the morphometric parameters of millions of osteocyte lacunae in human bone samples, using the previously validated 3D lacunar metrics of Mader et al. [18]. The rigor of the method is confirmed by accurate lacunar identification, a standard precision study [35], and the sensitive detection of differences between cortical and trabecular regions. The demonstrated validation highlights the value of the imaging method, and we believe this study will provide a rigorous foundation for future large-scale lacunar investigations.

## 2. METHODS

### 2.1. Human bone biopsy preparation

Thirty-one transiliac bone biopsy samples from premenopausal women already described in previous studies by Cohen et al. [36, 37] were used for this study. Biopsy samples had been obtained in women, aged 18-48, recruited as a reference population for studies of bone structure and metabolism in premenopausal women. Reference population subjects were required to have normal areal spine, hip, and forearm BMD by dual-energy x-ray absorptiometry (DXA; Z score ≥ −1.0 at all sites), no history of adult low trauma fracture, and no historical or biochemical evidence of diseases or conditions known to affect skeletal integrity [36, 37]. All subjects provided written informed consent; studies had been approved by the institutional review boards of all participating institutions.

A hemi-cylinder fraction of each biopsy core, containing both cortical and trabecular regions, was embedded in individual polymethylmethacrylate (PMMA) disks with a 25mm diameter. Subsections of each sample were prepared to fit in the desktop micro-CT scanner, which limits the diameter to a 4.0mm field of view (FOV) in the ultra-high-resolution mode. Hence, each PMMA embedded core was cut three times parallel to the longitudinal axis of the biopsy using a circular diamond blade (SCAN-DIA Minicut 40, SCAN-DIA GmbH & Co. KG, Germany) and custom-made mounts for sample fixation. This produced a rectangular block, which had the XY target dimensions of 4.25 +/- 0.1mm with the Z dimension depending on the original placement in the PMMA disk and ranging from 20-25mm. Each block was then turned on a conventional lathe (Schaublin 102, Bevilard, Switzerland) to create a final cylinder of 3.8 +/- 0.05mm diameter and a length ranging between 10 and 15mm. The biopsy cylinder was then tightly fit into a custom sample holder as seen in Figure 1 to minimize motion artifacts and to maximize the volume scannable within the 4.0mm diameter FOV.

**Figure 1:**
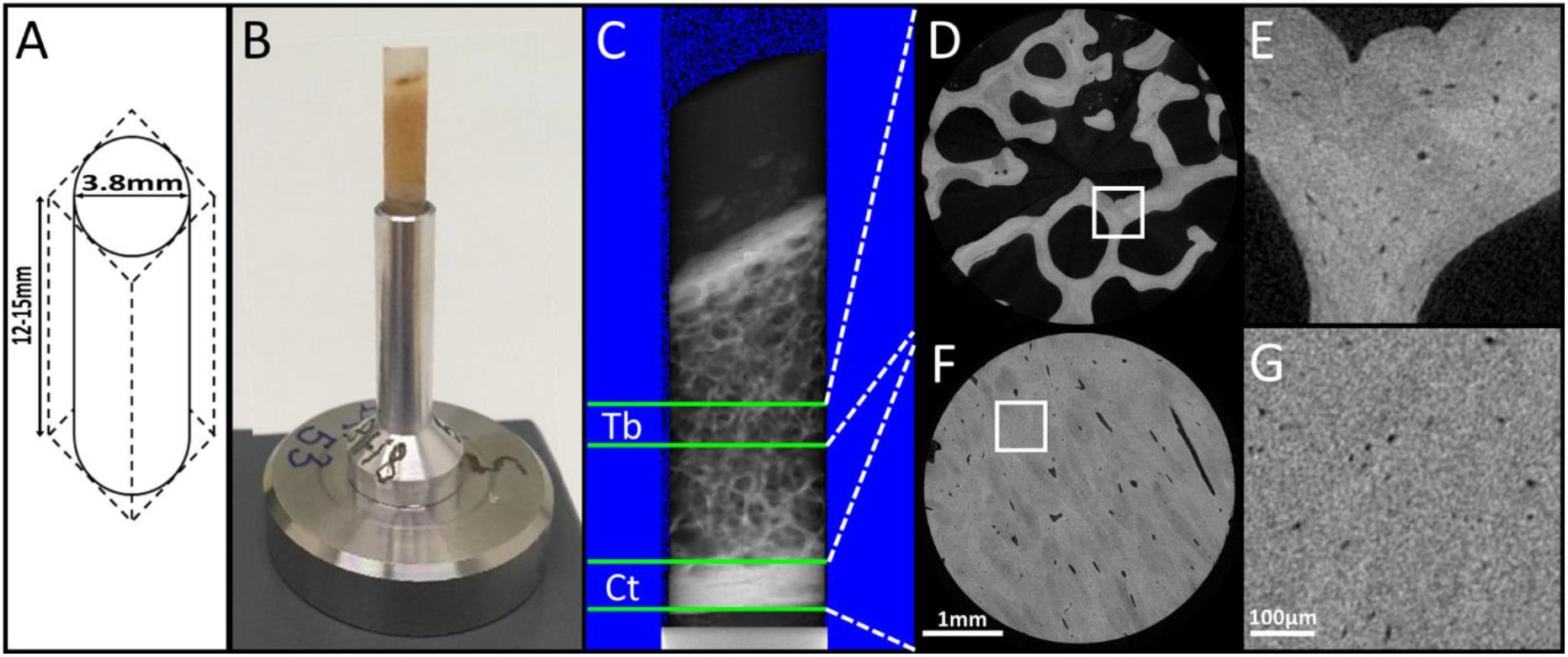
A) Schematic of sample core extraction from hemi-cylinder biopsy, which consisted of three linear cuts followed by a lathe turn. B) Photograph of final machined biopsy core inserted into a tolerance-fit sample holder. C) Scout-view overview of entire sample (XZ plane) with the trabecular (Tb) and cortical (Ct) scanned regions identified between the respectively labeled green lines. D) Ultra-high-resolution micro-CTscan of trabecular region cross-section (XY plane) and E) enlarged trabecular subregion. F) Ultra-high-resolution microCT scan of cortical region cross-section (XY plane) and G) enlarged cortical subregion.

### 2.2. Image Acquisition

Biopsy subsamples were imaged with a μCT50 (Scanco Medical AG, Brüttisellen, Switzerland), operated with a 0.5mm aluminum filter, 72μA current, 4W power, 55kVp energy, 1.5s integration time, level 6 data averaging, and with a total of 1500 projections. Images were reconstructed at a nominal isotropic voxel resolution of 1.2μm with an anti-ring level 8 to minimize center ring artifacts using the manufacturer’s scanner software. Each image consisted of a cylindrical volume equal to the full diameter of the sample (3.8 +/- 0.05mm) and the height of one scan stack (909 slices = 1.09mm). The protocol for each sample consisted of three scans: 1. Prescan to warm the sample in the scanner gantry in an effort to reduce motion artifacts caused by thermal effects (1 hour) 2. Cortical region scan starting from the lowest point on the centerline of the sample and scanning up one stack (10 hours) 3. Trabecular region scan stack in the middle of the biopsy equidistant between both cortical walls (10 hours). A visual example of this scanning protocol and resulting images can be seen in Figure 1.

To determine the optimal beam energy, three samples of trabecular bone were scanned at three beam energies: 55, 70, and 90kVp. These samples used for beam energy optimization originated from a separate study [38]; however, they were also human bone biopsies from the iliac crest and were considered to be comparable with our biopsy group. We aimed to maximize the image signal-to-noise ratio (SNR) just as previous studies relating to other anatomical bone sites had done [39]. The linear attenuation coefficient (raw signal) was measured for ten two-dimensional subregions (~0.25mm^2^) at every beam energy in three samples for 90 regions in total. SNR was then calculated by adapting the Firbank equation to account for two distinct materials as described in Equation 1 where μ is the average coefficient of linear attenuation of bone and the background (PMMA) and σ is the standard deviation of the background [40].

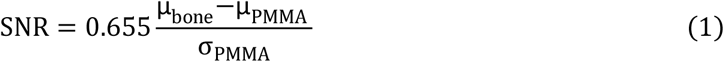

### 2.3. Image Preprocessing

Preprocessing of each image consisted of a constrained 3D Gaussian low pass filter (σ=0.8, support =1) to reduce noise and was applied using IPL (Scanco Medical AG, Brüttisellen, Switzerland). Segmentation of lacunar structures was performed by inverting the image after applying a threshold that was individualized for each image volume, which was necessary due to the large variation in tissue mineral density (TMD) distributions between samples at this resolution. The lacunar threshold was determined by fitting a Gaussian distribution to each sample’s raw TMD distribution data and calculating the first critical point (g’) of the fitted distribution using a custom Python script (3.7.1, Python Software Foundation, Delaware, USA). Bone volume (BV) was determined by fixing a threshold of 520mg HA/ccm, applying to all samples, performing a closing operation, and calculating the resulting BV using IPL software (Scanco Medical AG, Brüttisellen, Switzerland).

**Figure 2:**
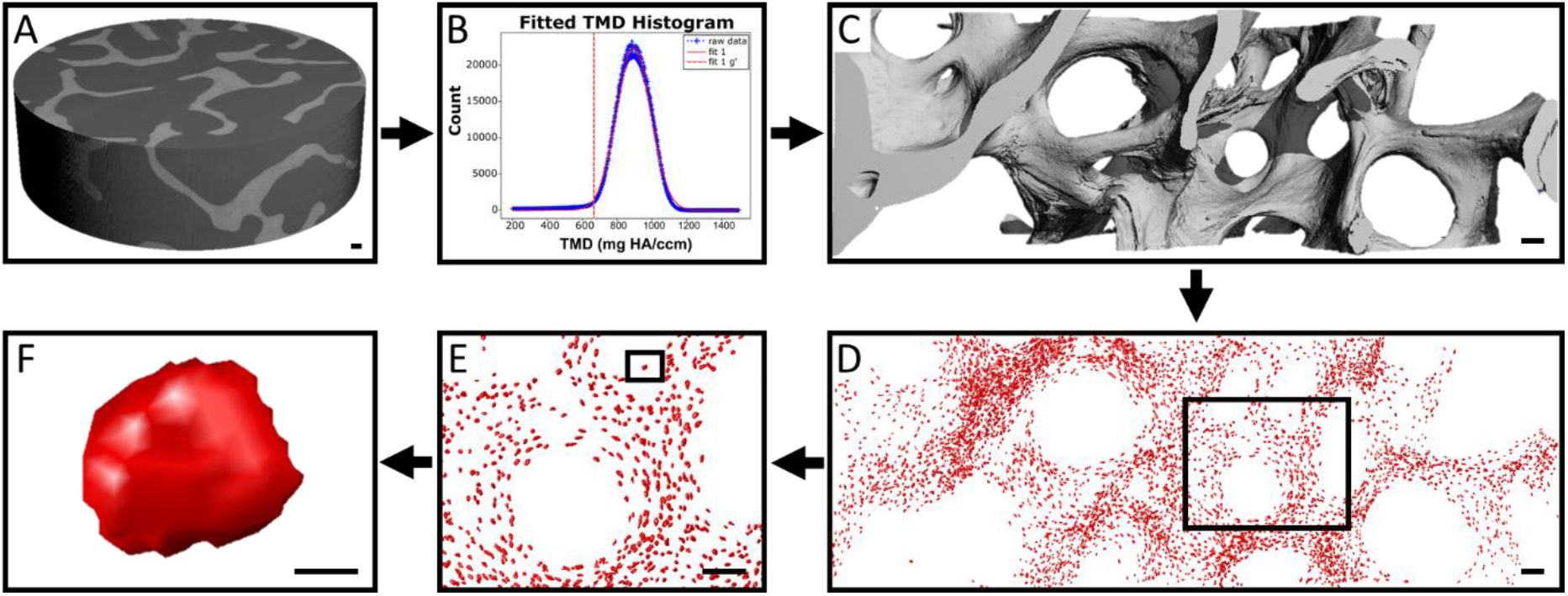
Visual overview of large-scale lacunar segmentation. A) Gaussian filtered, unsegmented, complete micro-CT image stack. B) Sample specific histogram of tissue mineral density (TMD) values fitted by a Gaussian function. A unique lacunar threshold is chosen for each sample at the first critical point of the respective fit by calculating the maximum of the first derivative (g’). C) Lacunar threshold calculated with (B) applied to (A). D) Image (C) inverted and component labeled to identify lacunae. E) Enlarged subregion group of lacunae. F) Single visualized lacuna. Scale bar lengths: A-E) 100μm F) 10μm.

All objects smaller than 50μm^3^ and larger than 2000μm^3^ were removed as to reflect the range of human lacunar volumes reported in previous histological studies [41]. The lower bound filters out noise structures while the upper bound excludes larger porosities like Haversian and Volkmann canals. Similar volumetric ranges have also been implemented in previous studies [42, 43]. Several microcracks, blood vessels, and image ring artifacts escaped the volumetric filter, yet all exhibited similar thin structures with a high object-elongation value. Therefore, these were excluded by removing all objects with an elongation above a threshold (Lc.St > 0.85). All objects sharing a border with the image edge were also removed to exclude partially cutoff objects.

### 2.4. Image Morphometry

The lacunar morphometries were calculated with a custom Python script, which first component labeled all lacunar objects, applied a surface mesh to each, and then measured basic individual parameters. Lacunar density (Lc.N/BV) was calculated by normalizing the number of lacunae (Lc.N) to the bone volume (BV) and lacunar porosity (Lc.TV/BV) by dividing the total volume of all lacunae by the respective BV. Local parameters (denoted with <> and first defined by Stauber et al. [44]) were each normalized to the sample Lc.N while population-based parameters (denoted with []) were not normalized. Individual lacunar volume (Lc.V), lacunar surface area (Lc.S), and the Eigen vectors were determined from the object specific mesh. This mesh was calculated by performing a triangulation of the surface voxels of the object using Lewiner marching cubes (3.7.1, Python Software Foundation, scikit-image library, Delaware, USA). The Eigen vectors were then used to quantify more complex parameters including lacunar stretch (Lc.St) and lacunar oblateness (Lc.Ob), which were first defined by Mader et al. [18]. Lacunar equancy (Lc.Eq) was the ratio between the smallest and largest Eigen vectors (E3/E1) [21, 45] while lacunar sphericity (Lc.Sr) related the lacunar object to a sphere via the ratio between Lc.S and Lc.V [43]. Lacunar angle (Lc.θ) was measured in degrees and ranged between 0 and 180 degree in relation to an arbitrarily created unit vector that was held consistent between images.

### 2.5. Validation

#### 2.5.1. Accuracy of automated lacunar identification

To evaluate the accuracy of the automated threshold approach, lacunae from five sample sub-volumes consisting of roughly 200 lacunae per region were manually identified by a trained observer by moving through stacks of slices and then compared with the automatically identified lacunae at the corresponding sample-specific g’ threshold to calculate true positive, false positive, and false negative rates (TPR, FPR, FNR). All objects were verified in 3D as depicted in Figure 3 and falsely classified objects were reclassified when appropriate. The geometry of false negatives was evaluated in 2D while the respective center voxel position of the objects could be evaluated in 3D in relation to successfully segmented true positive and false positive objects. Objects which were characterized as false positives typically were jagged, irregular in shape, and did not generally resemble lacunae in both 2D and 3D.

**Figure 3:**
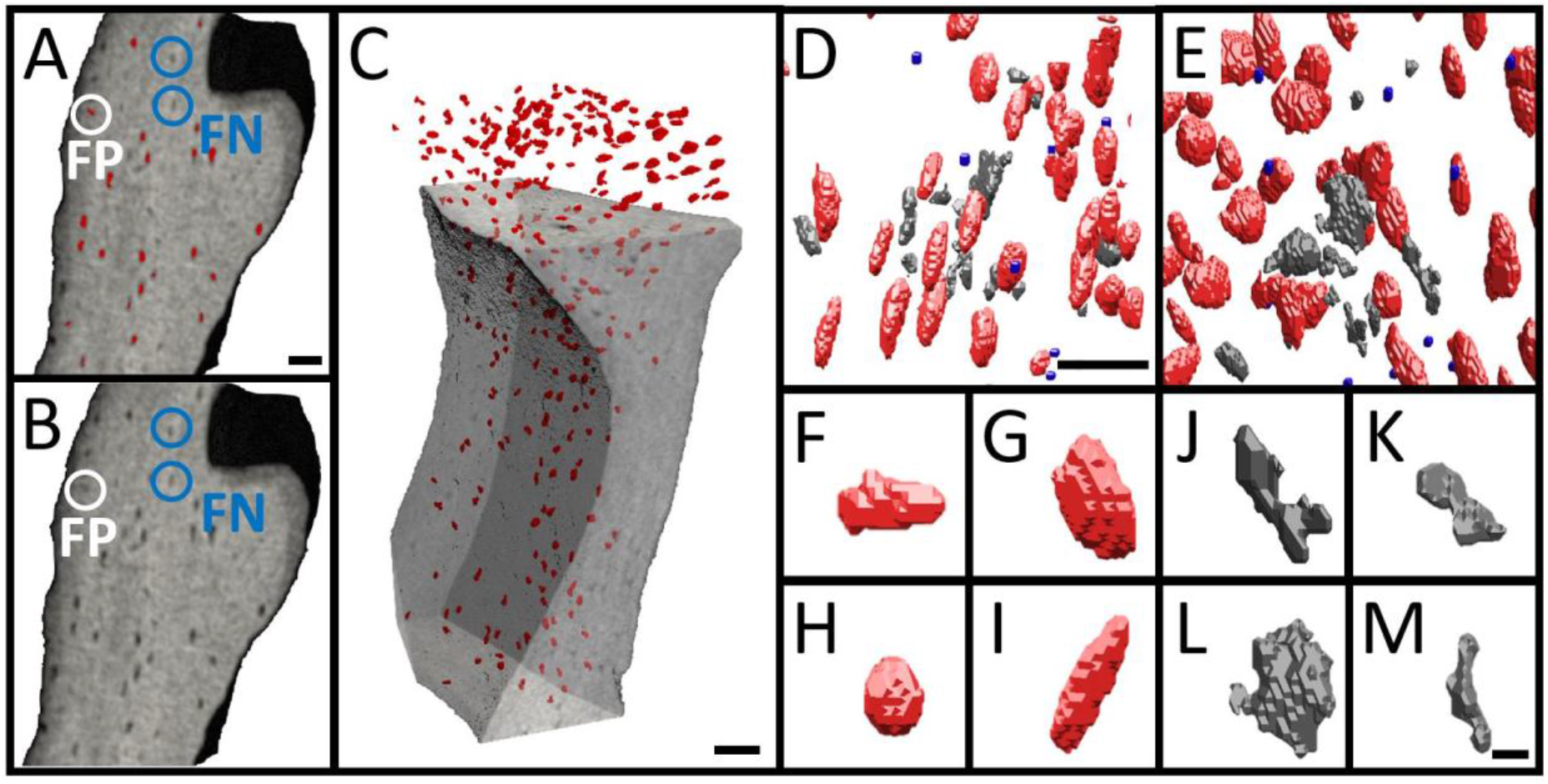
Manual vs. automatic lacunar identification. A) 3D cutplane of trabecular subregion with automatically segmented objects highlighted in red. Examples of false negatives (FN) circled in blue, false positives (FP) circled in white. B) Same cutplane as (A) used for visual comparison regarding classification of automatically segmented objects, showing same FP and FN as in (A). C) 3D orthographic projection of (A&B) with automatically segmented objects in red. D&E) 3D visualization comparing manual and automatic lacunar identification. Red objects = true positives (TP); objects identified as lacunae both manually and automatically. Gray objects = false positives (FP); objects identified as lacunae automatically but rejected manually. Blue spheres = false negatives (FN); center voxel of objects manually identified as lacunae but rejected by the automatic method. F-I) TP examples. J-M) FP examples. Scalebar A-E) = 50μm; F-M) = 10μm.

#### 2.5.2. Reproducibility of repeated measurements

In accordance with repeated measurements literature that recommends a sufficient number of degrees of freedom (DOF) to produce an upper confidence limit of the precision error that is 40% greater than the mean precision error [35], six samples were measured five times and repositioned between each measurement for a total of 20 DOF. Reproducibility of repeated measurements was evaluated by calculating the precision error (PE_%CV_) using Equation 2 – 3 where m is the quantity of subjects, SD represents the standard deviation of m repeated measures on subject j, and CV is the coefficient of variation. The intraclass correlation coefficient (ICC) for the measured lacunar indices was calculated using Equation 4 where n represents the number of repetitions and F0 represents the ratio of the residual within-subject mean squares and the between-subject mean squares.

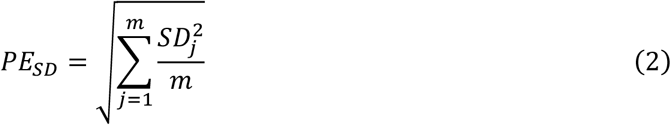

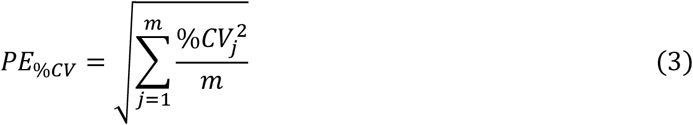

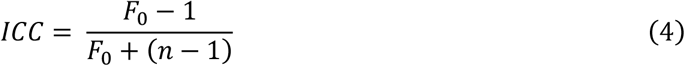

#### 2.5.3. Biological sensitivity of lacunar parameters

To assess the method’s sensitivity to biological differences, lacunar morphometric parameters from cortical and trabecular regions were compared as there are known physiological differences between the distribution and shape of lacunae in these regions in humans [43, 46]. Because each individual biopsy contained both cortical and trabecular regions, it was possible to compare lacunae both within samples and between samples. This biological sensitivity analysis was inspired by the previous work of Nebuloni et al. [47].

### 2.6. Statistical Analysis

A paired t-test with the necessary Bonferroni correction was performed with respect to energy-dependent imaging parameters in Table 1. With respect to the precision test, the 95% confidence interval was calculated for each morphometric parameter using a chi-squared distribution to gain an understanding of certainty with respect to the PE and ICC reported values in Table 3. Creating two-parameter plots of the data presented in Table 3 is another way to evaluate the reproducibility of the imaging method by means of clustering as is depicted in Figure 5. A paired Student’s t-test was performed to evaluate regional differences between lacunar parameters that were normalized to tissue indices (p<0.001). Population-based parameters were not normally distributed following a Kolmogorov-Smirnov test, and so a non-parametric Mann-Whitney U test was performed to evaluate population differences (p<0.001). The inter-quartile ranges and medians were computed for each morphometric parameter.

**Table 1:** Energy dependency of imaging parameters. Energy level 55kVp used as baseline and paired t-test with the necessary Bonferroni correction used to test for significant differences between energy levels in the three following categories: contrast, standard deviation of the image background (σ_PMMA_), and signal-to-noise ratio (SNR). Significantly different (p<0.005) from baseline denoted with (*) and from 70kVp with (#).

| Energy (kVp) | Contrast (n=30) | $\sigma_{\text{PMMA}}$ (n=30) | SNR (n=30) |
| --- | --- | --- | --- |
| 55 | 1.84±0.46 | 0.14±0.02 | 8.34±2.20 |
| 70 | 1.33±0.33* | 0.11±0.01* | 7.65±1.88 |
| 90 | 0.92±0.23*# | 0.09±0.01*# | 6.67±1.60* |

## 3. RESULTS

### 3.1. Image Acquisition

Images captured with a 55kVp beam energy exhibited a significantly higher SNR when compared to images obtained with a beam energy of 90kVp (p<0.005). The SNR measured with the 70kVp beam energy fell in between the high and low beam energy values and did not differ significantly from the other energies.

As shown in Table 1, both contrast and standard deviation of the background (PMMA in this case) were inversely proportional to the beam energy. Specifically, the inverse proportionality was approximately linear with contrast while σpMMA was quadratic (Figure 4), both important for the computation of SNR as given in Equation 1. As a ratio of both noise and the standard deviation of the image background (Equation 1), SNR was less dramatically affected by increasing beam energy. We therefore set the beam energy to 55kVp for all scans in the study because this setting produced the highest quality images with the highest contrast at acceptable noise levels.

**Figure 4:**
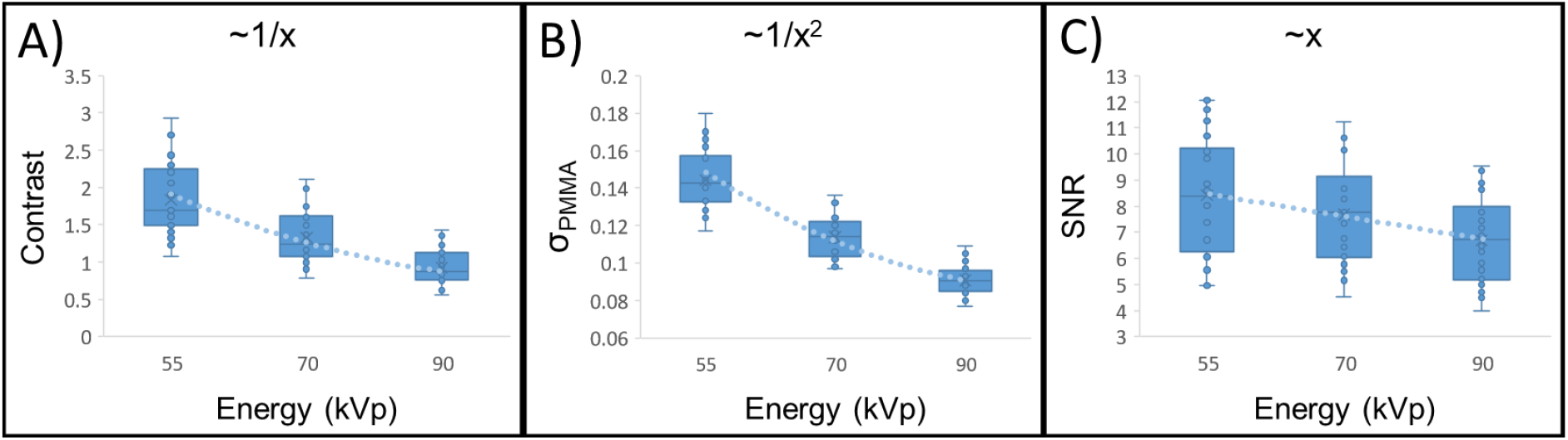
Beam physics relationships. Ten subregions measured at each energy (30 measurements in total). A) Contrast exhibits an inverse relationship with beam energy. B) Standard deviation of the background (σ_PMMA_) has an approximate quadratic relationship with beam energy. C) Signal-to-noise ratio is linearly related to beam energy as expected since it is defined as the contrast divided by the σ_PMMA_.

### 3.2. Validation

#### 3.2.1. Accuracy of automated lacunar identification

Automatically identified objects were compared with the manually identified objects to evaluate the true positive rate (TPR), false positive rate (FPR), and false negative rate (FNR) measures as seen in Table 2. Objects identified automatically in each sample revealed a strong agreement with the lacunae counted manually as is evident by the high TPR and low FPR and low FNR. Roughly 200 lacunae were present in each sample subregion with sample 1 exhibiting the highest TPR (93.9%) and lowest FNR (6.1%). Sample 4 had the lowest FPR (0.5%) while sample 3 the lowest TPR (77.3%). The final lacunar identification accuracy measure was calculated as the mean of the five samples and was computed to be 89.0% TPR, 3.4% FPR, and 11.0% FNR.

**Table 2:**
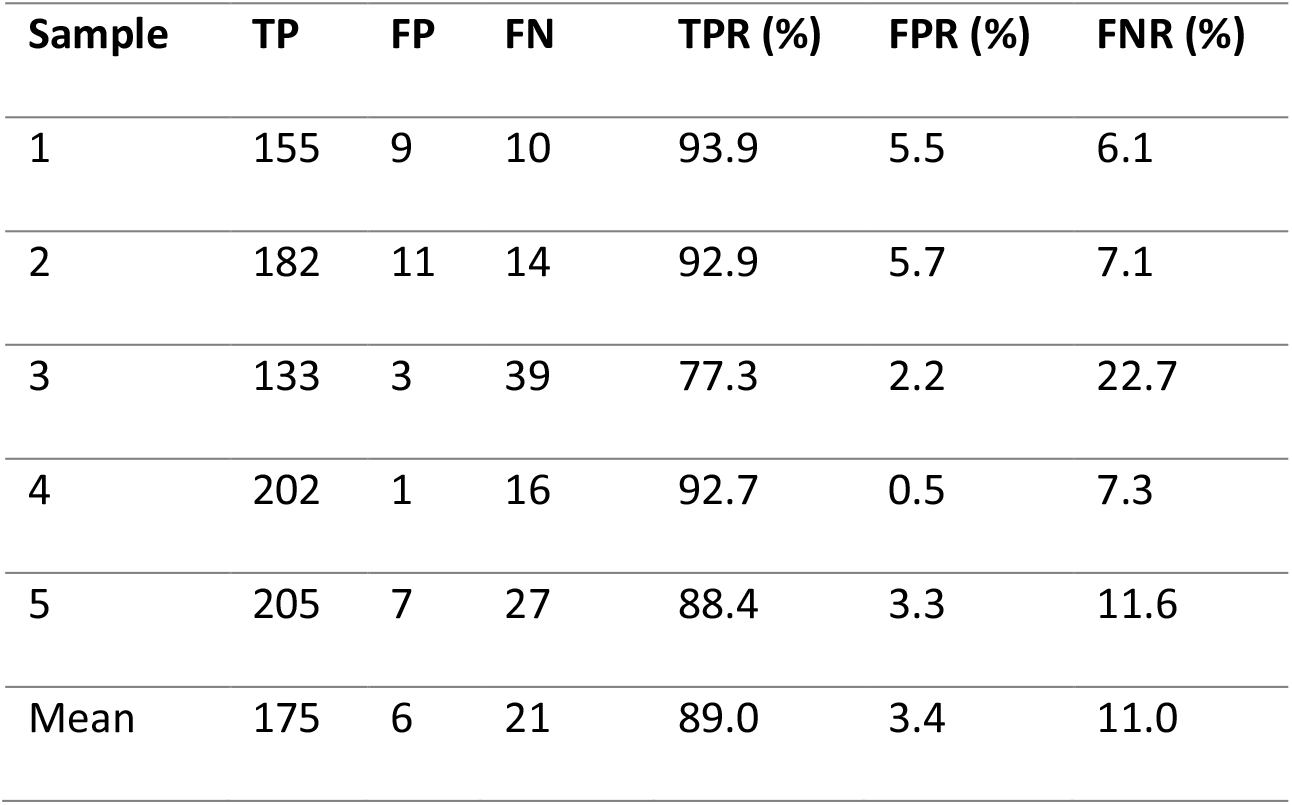
Quantification of accuracy measures and their corresponding rates obtained via comparison between manual and automatic lacunar identification methods. TPR = True Positive Rate; FPR = False Positive Rate; FNR = False Negative Rate.

**Table 3:** Reported values from the reproducibility analysis (n=6, five repeated measurements). Morphometric parameters include: lacunar total volume (Lc.TV), lacunar porosity (Lc.TV/BV), lacunar number (Lc.N), lacunar density (Lc.N/BV), local lacunar volume (<Lc.V>), local lacunar surface area (<Lc.S>), local lacunar stretch (<Lc.St>), local lacunar oblateness (<Lc.Ob>), local lacunar sphericity (<Lc.Sr>), local lacunar equancy (<Lc.Eq>), and local lacunar angle (<Lc.θ>). In addition to mean values of each morphometric parameter across the samples, we report the following for precision errors (PE): standard deviation (PE_SD_) in absolute values, coefficient of variation (PE_%CV_) of the repeated experiments, and the 95% confidence interval of the variation (95% CI PE%CV). Also reported are the intraclass correlation coefficients (ICC) and the 95% confidence interval of the ICC for each respective morphometric parameter.

| **Morphometric parameter** | **Mean** | **PE<sub>SD</sub>** | **PE<sub>%CV</sub>** | **95% CI PE<sub>%CV</sub>** | **ICC** | **95% CI ICC** |
| --- | --- | --- | --- | --- | --- | --- |
| Lc.TV (1000*μm<sup>3</sup>) | 2,119.3 | 40.2 | 2.77% | 2.11-4.05% | 0.997 | 0.992-1.000 |
| Lc.TV/BV (%) | 0.5 | 0.009 | 2.52% | 1.91-3.68% | 0.989 | 0.966-0.998 |
| Lc.N (1000) | 10.03 | 0.2 | 1.86% | 1.41-2.72% | 0.999 | 0.996-1.000 |
| Lc.N/BV (1000/mm<sup>3</sup>) | 22.09 | 0.3 | 1.48% | 1.12-2.16% | 0.984 | 0.952-0.997 |
| <Lc.V> (μm<sup>3</sup>) | 210.5 | 2.5 | 1.33% | 1.01-1.95% | 0.991 | 0.970-0.999 |
| <Lc.S> (μm<sup>2</sup>) | 207.5 | 1.7 | 0.88% | 0.67-1.29% | 0.990 | 0.968-0.998 |
| <Lc.St> (1) | 0.6 | 0.001 | 0.17% | 0.13-0.25% | 0.989 | 0.963-0.998 |
| <Lc.Ob> (1) | -0.4 | 0.006 | 1.59% | 1.21-2.32% | 0.960 | 0.883-0.994 |
| <Lc.Sr> (1) | 0.8 | 0.002 | 0.24% | 0.18-0.35% | 0.938 | 0.816-0.990 |
| <Lc.Eq> (1) | 0.3 | 0.001 | 0.43% | 0.33-0.63% | 0.994 | 0.980-0.999 |
| <Lc.θ> (Degree) | 113.8 | 0.8 | 0.66% | 0.50-0.97% | 0.980 | 0.938-0.997 |

#### 3.2.2. Reproducibility of repeated measurements

Repeated measures reproducibility is crucial for validation and was quantified for lacunar indices. Across all lacunar morphometric parameters, precision errors were very low (below 3%) and the ICC were very high (above 0.980), which indicate that the measurements of these lacunar parameters are extremely reproducible. The 95% confidence interval range for precision errors was between 0.67% and 3.68% while the range for ICC values were between 0.883 and 1.000, which indicates extremely low measurement variability and high reproducibility.

Clustering of the individual repeated measurements indicates reproducibility of lacunar morphometry and is especially apparent in Figure 5A-B. These parameters also exhibited a very high correlation, which illustrated that as bone volume increases so will the number of lacunae and the total lacunar volume. The clustering in Figure 5C-D was not nearly as evident across all six samples due to the fact that <Lc.V> and <Lc.St> were more difficult to reproduce. As BV/TV increases in Figure 5C-D, measured variability decreases between the first and second cluster and then remains approximately constant. The strong correlation of Lc.N (R^2^ = 0.99) and Lc.TV (R^2^ = 0.98) with BV/TV position them as potential lacunar biomarker candidates. <Lc.V> and <Lc.St> were more independent of BV/TV and no correlation was found.

**Figure 5:**
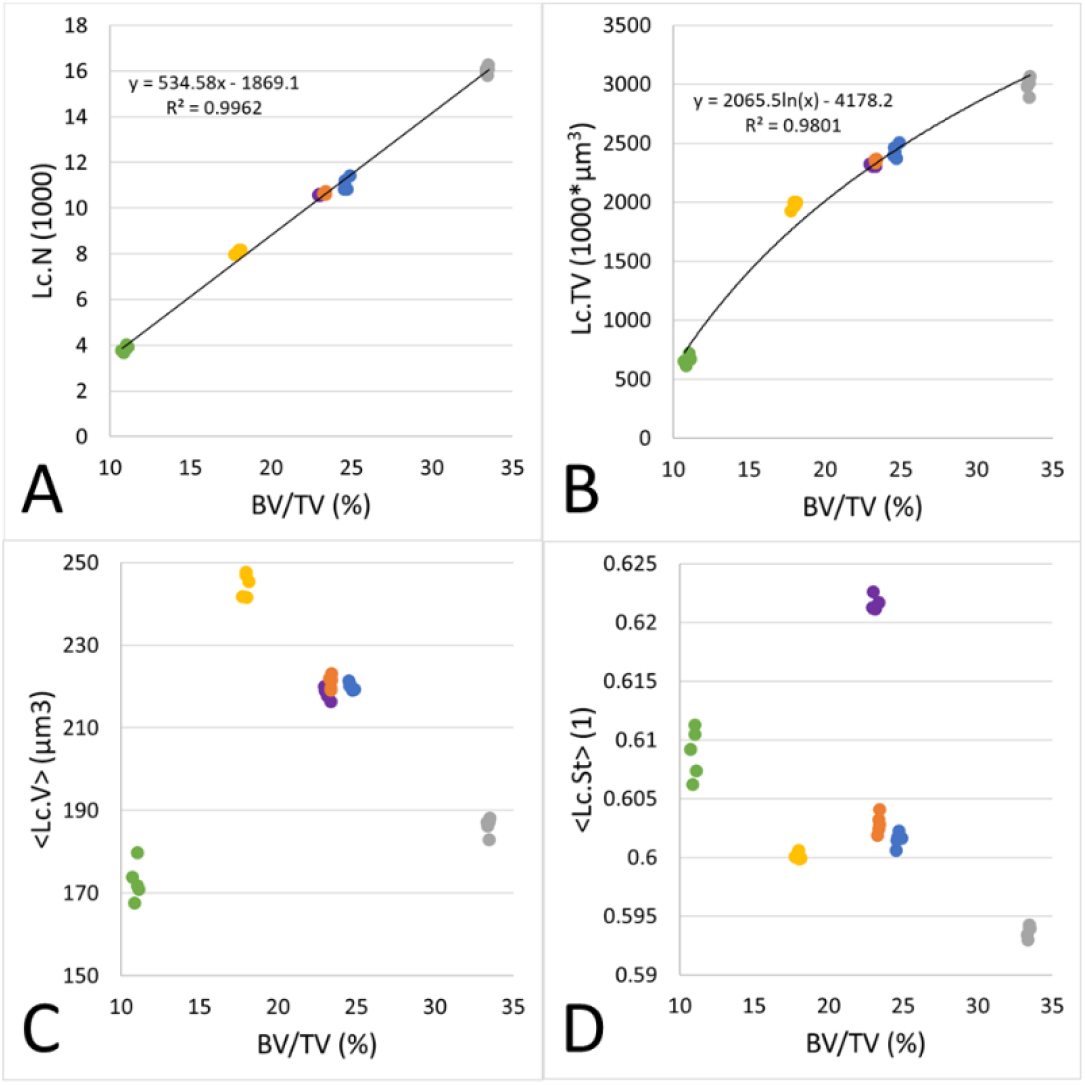
Two-parameter plots that demonstrate the reproducibility of the imaging method. Each color represents an individual sample and each data point represents a specific measurement (n=6, with 5 repeated measurements). A) Lacunar number (Lc.N) vs. bone volume (BV/TV). B) Lacunar total volume (Lc.TV) vs. bone volume (BV/TV). C) Local lacunar volume (<Lc.V>) vs. bone volume (BV/TV). D) Local lacunar stretch (<Lc.St>) vs. bone volume (BV/TV).

#### 3.2.3. Biological sensitivity of lacunar parameters

Cortical and trabecular bone regions were measured and compared to evaluate the biological sensitivity of the method. More bone was present, and consequently, more lacunae were present in cortical bone when compared to trabecular bone. Measured tissue values such as BV, and BV/TV in trabecular regions were consistently lower than cortical regions as expected since trabecular bone is sparse and cortical bone is compact. Regarding the global morphometries, which were normally distributed, Lc.N/BV median value in trabecular bone was nearly half of what it was observed to be in cortical bone (16,611 vs. 26,429, p<0.001). Similarly, the median value of Lc.TV/BV in trabecular bone was also nearly half of what it was in cortical bone (0.30% vs. 0.58%, p<0.001), again indicating that cortical bone has a significantly higher lacunar porosity than trabecular bone. Furthermore, we report in Table 4 that the normalized local parameters <Lc.V> and <Lc.S> are significantly greater (p<0.001) in cortical bone (<Lc.V> = 223μm^3^; <Lc.S> = 233μm^2^) than in trabecular bone (<Lc.V> = 178μm^3^; <Lc.S> = 194μm^2^). Most population-based morphometric parameters were not normally distributed, and consequently we reported the median values as well as the interquartile range for all indices to provide a sense of the distribution of each parameter in Table 4.

**Table 4:** Large-scale lacunar morphometric parameters (n=6.57 million for cortical and n=1.14 million for trabecular). Reported values are the median with interquartile range (25^th^ percentile – 75^th^ percentile). Morphometric parameters include: total image volume (TV), total bone volume (BV), ratio of bone volume to total volume (BV/TV), lacunar porosity (Lc.TV/BV), lacunar number (Lc.N), lacunar density (Lc.N/BV), local lacunar volume (<Lc.V>), local lacunar surface area (<Lc.S>), population lacunar volume ([Lc.V]), population lacunar surface area ([Lc.S]), population lacunar surface area to volume ratio ([Lc.S/Lc.V]), population lacunar stretch ([Lc.St]), population lacunar oblateness ([Lc.Ob]), population lacunar sphericity ([Lc.Sr]), population lacunar equancy ([Lc.Eq]), and population lacunar angle ([Lc.θ]). Paired t-test performed for the normally distributed global and local parameters, (*) indicates p<0.001. Mann-Whitney U test performed on populationbased parameters and (*) indicates p<0.001.

| Morphometric parameter | Trabecular | Cortical |
| --- | --- | --- |
| TV (mm<sup>3</sup>) | 15.7 (15.1–15.8) | 15.7 (15.6–15.9) |
| BV (mm<sup>3</sup>) | 2.11 (1.54–2.46)* | 7.68 (6.33–9.70) |
| BV/TV (%) | 14.0 (10.2–18.4)* | 57.8 (43.7–64.9) |
| Lc.TV/BV (%) | 0.30 (0.22–0.38)* | 0.58 (0.51–0.65) |
| Lc.N (1000) | 41.2 (27.7–67.9)* | 252 (175–266) |
| Lc.N/BV (1000/mm<sup>3</sup>) | 16.6 (14.1–18.7)* | 26.4 (23.7–29.3) |
| $\langle \text{Lc.V} \rangle$ (μm<sup>3</sup>) | 178 (159–197)* | 223 (189–245) |
| $\langle \text{Lc.S} \rangle$ (μm<sup>2</sup>) | 194 (178–211)* | 233 (216–248) |
| [Lc.V] (μm<sup>3</sup>) | 123 (70.1–230)* | 116 (63.0–272) |
| [Lc.S] (μm<sup>2</sup>) | 161 (109–245)* | 164 (105–290) |
| [Lc.S]/[Lc.V] (1/μm) | 1.30 (1.05–1.56)* | 1.41 (1.06–1.66) |
| [Lc.St] (1) | 0.62 (0.53–0.69)* | 0.61 (0.51–0.69) |
| [Lc.Ob] (1) | -0.44 (-0.67– -0.13)* | -0.49 (-0.71– -0.17) |
| [Lc.Sr] (1) | 0.76 (0.70–0.81)* | 0.73 (0.66–0.79) |
| [Lc.Eq] (1) | 0.29 (0.19–0.45)* | 0.31 (0.19–0.47) |
| [Lc.θ] (Degree) | 104 (76.0–131)* | 107 (77.4–134) |

Population-based lacunar parameters were not normally distributed. The interquartile ranges between regions were similar, yet a non-parametric Mann-Whitney U test revealed significant differences between cortical and trabecular regions (p<0.001). The measure of sphericity ([Lc.Sr]) was approximately 0.75 for both regions of bone, indicating similarities between the measured ellipsoids and an idealized sphere. The additional parameters [Lc.Eq], [Lc.St] and [Lc.Ob] allow for the ellipsoidal lacunar shape to be described in more detail, and the median reported values suggest that these are indeed ellipsoidal structures in both trabecular and cortical regions. The range of [Lc.Ob] in trabecular bone was slightly higher than in cortical bone.

Figure 6 depicts selected global, local, and population-based morphometries from both cortical and trabecular regions. When normalized to the analyzed tissue volume, both the lacunar density and porosity were significantly different between cortical and trabecular regions (Figure 6A-B) across all 31 samples, further supporting their potential as biomarkers. Local morphometries (<Lc.V> & <Lc.S>) were also significantly different between the two regions (Figure 6C-D), yet not as clearly separated as the global morphometries. Population-based morphometries (Figure 6E-F) included all lacunar observations across all 31 samples: 1.14 million lacunae in trabecular bone and 6.57 million lacunae in cortical bone. The shape indices [Lc.St] and [Lc.Sr] were chosen to compare between regions and were also significantly different. A visual comparison is presented in Figure 6G-H between the samples containing the median Lc.N/BV values from Figure 6A, further illustrating the differences between anatomical regions.

**Figure 6:**
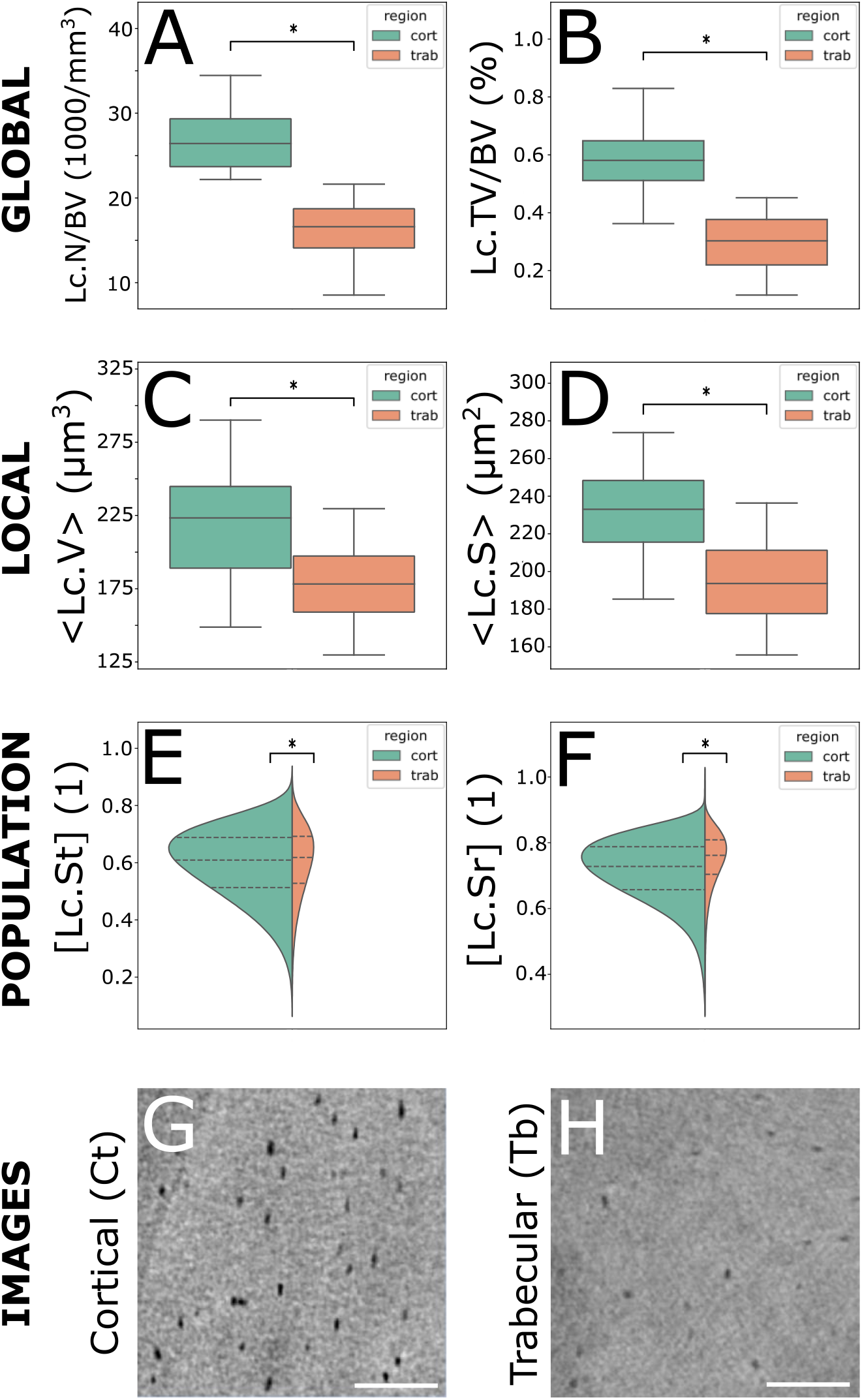
Comparison of cortical and trabecular regions regarding lacunar morphometric parameters. Included in the analysis were 31 samples which comprised of 6.57 million cortical lacunae and 1.14 million trabecular lacunae. A) Lacunar density (Lc.N/BV). B) Lacunar porosity (Lc.TV/BV). C) Local lacunar volume (<Lc.V>) normalized to sample. D) Local lacunar surface area (<Lc.S>) normalized to sample. E) Population-based local lacunar stretch ([Lc.St]) not normalized to sample. F) Population-based local lacunar sphericity ([Lc.Sr]) not normalized to sample. G&H) Micro-CT images from cortical and trabecular regions representing the median values from plot (A) respectively, scale bar = 100μm. Paired t-test performed on normalized parameter plots (AD) and (*) indicates p<0.001. Mann-Whitney U test performed on population-based parameters (E-F) and (*) indicates p<0.001.

### 4. DISCUSSION

Detailed examination of the LCN on a large scale demands a high-resolution 3D-imaging methodology that is accurate, reproducible, and sensitive. Lacunae must be identified accurately and morphometric indices measured repeatably. Therefore, it is paramount that researchers select a validated imaging methodology for the large-scale assessment of osteocyte lacunar morphometry. High-resolution desktop micro-CT is an ideal technology for large-scale investigation of the lacunar network due to its wide accessibility. Micro-CT has been employed for decades as a standard technology for bone tissue morphometry with nominal voxel resolutions in the range of 10-40μm [25–28]. However, this technology has evolved in recent years, and with it, the ability to image higher resolutions on the order of 1μm [48]. Hence, the development of validated image acquisition, processing, and analysis tools to image lacunar morphometry is a logical progression of the science as well as being crucial to understanding the biological impact of the LCN on bone at other hierarchical levels.

Identifying the lacunar structures to be analyzed required a landmark-based threshold to be applied to the image that accounted for sample-specific mineralization heterogeneities. Typically, a single threshold is chosen by the user via visual inspection and then applied to all samples in a study. At the tissue level this method is acceptable when recommended guidelines are carefully followed [49] and was how BV was calculated in this study. However, due to the wide variation of the TMD distributions between samples at the 1.2μm resolution, this single-threshold method cannot be applied to all samples when imaging lacunae (Figure S1). Therefore, our approach was to pragmatically locate a threshold that was intrinsically linked to the sample specific TMD distribution, compare the resulting lacunar identification with manual identification, and quantify the accuracy. Previous studies have tried similar individual threshold approaches, which are offset from a reference point of the TMD histogram such as the mean [50], but we found our specific images responded best to selecting the TMD histogram critical point as a landmark for segmentation (Figure S1). In addition, the critical point is a clear and calculable landmark on every TMD histogram, which is less prone to error than an offset. This produced an adequate automated segmentation of lacunar structures as is depicted in Figure 3. Alternative descriptors of the TMD distribution were investigated such as the width of the distribution but did not prove to be as effective as the first critical point (Figure S2). It is important to note that the TMD distribution can be very heterogeneous around osteocyte lacunae and that adjustments to the algorithm would need to be made when investigating topics like osteocytic osteolysis which changes the mineral surrounding the lacunar structures. The μCT50 machine was calibrated weekly with a multi-material phantom to minimize measurement errors with respect to scaling the raw signal attenuation to values of hydroxyapatite for the TMD histograms.

The range of considered object volumes was also important for lacunar identification. This range varies substantially between studies and could be as narrow as 50-610μm^3^ or as wide as 175-2000μm^3^ [18, 21, 42, 43, 51, 52]. Previous examinations of histological slides have estimated the human lacunae to be between 28μm^3^ and 1713μm^3^, yet lacunae observed below 50μm^3^ were only found in fracture callus [41]. After evaluating several different volume ranges and comparing both qualitatively with 2D and 3D images and quantitatively with accuracy measures such as TPR, FPR, and FNR, we determined a lower limit of 50μm^3^ to be optimal for lacunar identification in human trabecular and cortical bone [53]. The upper limit was chosen to be 2000μm^3^ in line with a similar study [21]. This range has a large impact on the number of lacunae analyzed and is particularly sensitive on the lower limit. This problem is especially pronounced with respect to desktop micro-CT due to the limited photon count of its X-ray beam technology. Relative to studies conducted with synchrotron CT systems [18, 42], the desktop micro-CT X-ray beam creates image projections with fewer photons, which increases the resulting noise, reduces the image quality, and makes visualization of small lacunae more difficult. A Gaussian filter with a low sigma value of 0.8 was applied so noise would be reduced while the borders of the lacunae would not be blurred beyond the recognition of a trained human observer. Several extremely elongated ring artifacts were observed in segmentations and were successfully removed by implementing an anti-ring reconstruction filter and applying a shape filter that removed objects with a Lc.St value greater than 0.85 [54].

We used ultra-high-resolution desktop micro-CT to image 7.71 million osteocyte lacunae across cortical and trabecular regions in 31 human iliac crest biopsies. We have observed that morphometric differences exist between lacunae in cortical and trabecular regions of bone, which has also been shown in previous studies [43, 46]. Currently, only Akhter et al. [43] have reported sample matched trabecular and cortical lacunar morphometric parameters in human iliac crest biopsies. In contrast to their study, we report higher values of Lc.N/BV and Lc.TV/BV in cortical bone when compared to trabecular bone. However, the narrow volume range they evaluate (50-610μm) and their analysis of less than 1% of the number of lacunae that we examine severely limits the range of variation that they could potentially consider. Additionally, we have proven our desktop micro-CT imaging technique to be accurate for lacunar identification, reproducible for repeated measures, and sensitive for lacunar parameters between anatomically distinct regions.

We evaluated not only global lacunar parameters related to tissue measures (Figure 6A-B), but also local (Figure 6C-D) and population-based (Figure 6E-F) values. Population-based morphometry was not normalized and presents the reader with an undistorted perspective of the natural variation of certain morphometric indices across millions of lacunae (Figure 6E-F). This further illustrates the method’s sensitivity that we see in Figures 6A-D and also demonstrates the method’s ability to capture the natural variation of lacunae in a large-scale analysis.

Manual identification was used as our gold-standard for calculating accuracy, because registration between micro-CT images and typical morphological gold-standards like histology is extremely difficult. However, we were careful to create the best manual identification dataset possible for comparison. Cresswell et al. have used similar accuracy comparisons in previous studies and in fact, achieve similar rates of TP, FP, and FN to ours [50]. Interestingly, sample 3 in our accuracy calculation exhibited an inordinately low TPR and high FNR. This was due to a suboptimal selection of the sample’s subregion near the bone surface, which made manual identification of lacunae slightly more difficult. FN objects could only be visualized in 2D as the machine incorrectly did not segment these objects. However, the 3D position of the center voxel in the FN objects could still be evaluated in relation to TP and FP objects as this was the coordinate of the voxel chosen by the trained observer. Objects determined to be FP resembled noise structures and were either image artifacts or non-lacunar micropores.

We observed very low precision errors and high intraclass correlation coefficients with respect to our five consecutive measurements of six samples. These values were in the same range as in the study of Hemmatian et al. who investigated the reproducibility of desktop micro-CT for imaging murine lacunae [48]. The tight clustering of data points when creating two-parameter plots as seen in Figure 5 further proves the reproducibility of the method. Figure 5A and 5B both exhibit tight clustering within the measurements for each sample which is what we expect when comparing lacunar parameters with tissue values like BV/TV. Bone tissue volume is a quantity that micro-CT is excellent at measuring and hence we would expect it to be extremely reproducible. Physiologically speaking, we would also expect Lc.N and Lc.TV to increase with increasing total bone volume, which explains the strong correlation, further adds credibility to the imaging modality, and even positions the two lacunar parameters as potential biomarker candidates. Previous research has demonstrated osteocytic osteolysis plays a role in bone health, disease, and mechanisms of medication response; this also poises the lacunar microarchitecture as a potential biomarker from the perspective of osteocyte-controlled bone remodeling [55]. Furthermore, a variety of chemically-based bone biomarkers, such as serum CTX and P1NP, currently exist for the clinical assessment of osteoporosis/bone remodeling and lacunar parameters like Lc.N and Lc.TV could potentially be used in conjunction with those recently outlined by Kuo et al. for investigation of disease mechanisms [56]. Yet in Figure 5C and 5D, we note that the values of each sample are slightly less clustered in comparison with Figure 5A and 5B and are not correlated. More specifically, we note that BV/TV remains very consistent but the <Lc.V> and <Lc.St> varies. Small objects are more difficult to mesh and both <Lc.V> and <Lc.St> are dependent on the object mesh which would explain the difficulties with reproduction in comparison to Lc.N and Lc.TV. Hemmatian et al. also found lacunar measures such as <Lc.V> to be less reproducible than tissue measures such as BV/TV [48]. Consequently, <Lc.V> and <Lc.St> do not appear to be good biomarker candidates.

Previous studies have demonstrated that lacunar morphometric parameters differ between trabecular and cortical regions of bone [43, 46, 57]. We used this observation to evaluate the biological sensitivity of our method by the ability to differentiate lacunae between anatomically distinct regions. This was based upon the analysis performed by Nebuloni et al. regarding the biological sensitivity of microCT relative to vascular imaging [47]. We report significant differences between global, local, and population-based parameters including Lc.TV/BV, Lc.N/BV, <Lc.V>, <Lc.S>, [Lc.St], and [Lc.Sr] as seen in Table 4 and Figures 6A-F. Furthermore, Figures 6G-H provide a visual confirmation of the difference that we report in Figure 6A. Not only do we see that lacunar density is lower in trabecular regions, but also the lacunae themselves look to be slightly smaller relative to the cortical regions. This would indicate that lacunae in trabecular regions also consist of lower total porosity (Lc.TV/BV), volume (<Lc.V>), and surface area (<Lc.S>). These visual differences further support our claim that the method is sensitive to previously studied regional differences and reflects the statistically significant differences that we report in Figures 6A-D. Kegelman et al. found similar lacunar morphometries in a recent study; however, overall their lacunae were slightly smaller, likely because they were of murine origin [57].

While we were able to evaluate the accuracy of lacunar identification between human and machine counting, we did not address the accuracy of the segmentation of the lacunar structures as other studies have done [48]. This is a limitation of our study and would be interesting to quantify. However, because lacunar segmentation is resolution dependent and Hemmatian et al. demonstrated that lacunar geometry is correlated, but not accurate, between desktop microCT and CLSM, we believe lacunar identification accuracy is more meaningful [48]. Additionally, the lacunar data were very sensitive to the selection of the lower volumetric bound. This was difficult to select since volumetric data from previous studies regarding the distinction between a lacuna and a micropore is limited. Furthermore, we must acknowledge the fact that partial volume effects generate error with respect to the segmented lacunar structures. Considering many lacunae have a diameter of roughly 10μm, these partial volume effects are accentuated at the 1.2μm voxel resolution. Finally, the precision study required weeks of scanning time and consequently was only performed on trabecular bone.

### 5. CONCLUSION

We present a new high-throughput method that is reproducible and sensitive to assess osteocyte lacunar morphometry in human bone samples. We use ultra-high-resolution desktop micro-CT, an individualized histogram-based segmentation procedure, and a custom evaluation algorithm to calculate global and local morphometric parameters of 7.71 million lacunae in two distinct regions of 31 human iliac crest bone samples, revealing two potential biomarkers. The validation of our method demonstrates accurate lacunar identification, reproducibility of repeated measurements, and sensitivity between anatomically distinct regions. Therefore, our new image acquisition and evaluation methodologies expand the number of investigable hypotheses surrounding osteocyte lacunae, while simultaneously employing a widely accessible and mature imaging technology – desktop micro-CT.

## ACKNOWLEDGEMENTS

The authors would like to thank the Joint Scoliosis Research Center of The Chinese University of Hong Kong and Nanjing University, Hong Kong & Nanjing, China for their joint efforts on providing and processing the bone biopsies used for beam energy optimization. We thank Lucid AG for assisting with the resolution dependency study using XamFlow. We would also like to thank Peter Schwilch for assisting with biopsy machining and Dr. Patrik Christen for his mentoring.

## ABBREVIATIONS

LCN: lacuno-canalicular network
FIB/SEM: focused ion beam/scanning electron microscopy
CLSM: confocal laser scanning microscopy
SR-CT: synchrotron radiation computed tomography
micro-CT: micro-computed tomography
DXA: dual-energy x-ray absorptiometry
BMD: bone mineral density
PMMA: polymethylmethacrylate
FOV: field of view
SNR: signal to noise ratio
IPL: image processing language software
TMD: tissue mineral density
BV: bone volume
BV/TV: bone volume / total volume
< >: denotes local lacunar parameter
[ ]: denotes population lacunar parameter
Lc.N/BV: lacunar density
Lc.N: lacunar number
Lc.TV: lacunar total volume
Lc.TV/BV: lacunar porosity
Lc.V: lacunar volume
Lc.S: lacunar surface area
Lc.St: lacunar stretch
Lc.Ob: lacunar oblateness
Lc.Eq: lacunar equancy
Lc.Sr: lacunar sphericity
Lc.θ: lacunar angle
TPR: true positive rate
FPR: false positive rate
FNR: false negative rate
DOF: degrees of freedom
PE: precision error
ICC: intraclass correlation coefficient
σ_PMMA_: standard deviation of the image background
CTX: C-terminal telopeptide of type 1 collagen
P1NP: Procollagen type I N-terminal propeptide

## SUPPLEMENTARY DOCUMENT

Below are several figures which provide additional information regarding the specifics of the imaging methodology.

**Figure S1:**
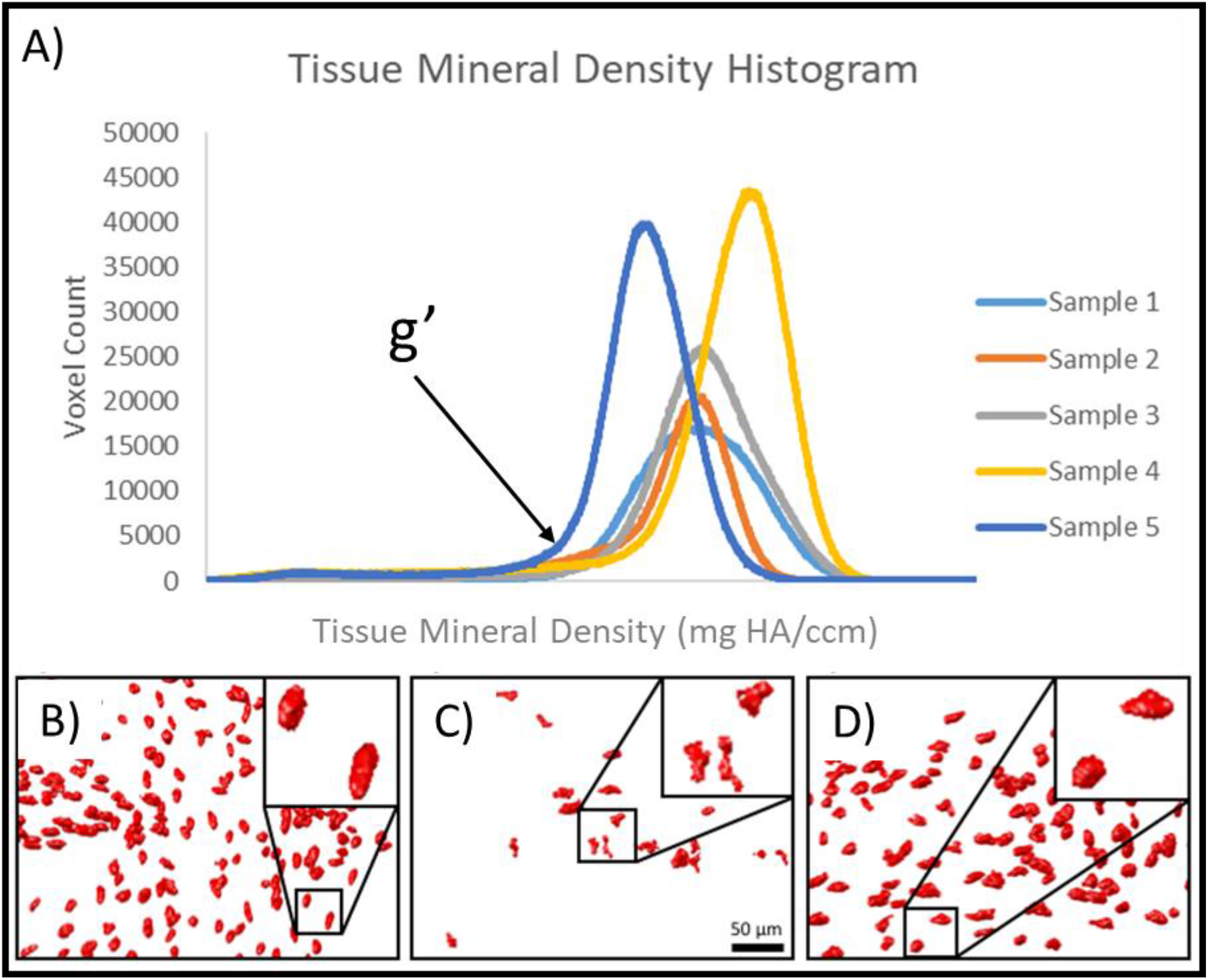
A) TMD histograms for several samples where g’ is the critical point. B) Segmented lacunae using a fixed threshold applied to sample 1. C) Segmented lacunae using the same fixed threshold that was applied to sample 1 to sample 2. D) Individualized threshold approach (g’) calculated for sample 2 and the resulting segmented lacunae.

**Figure S2:**
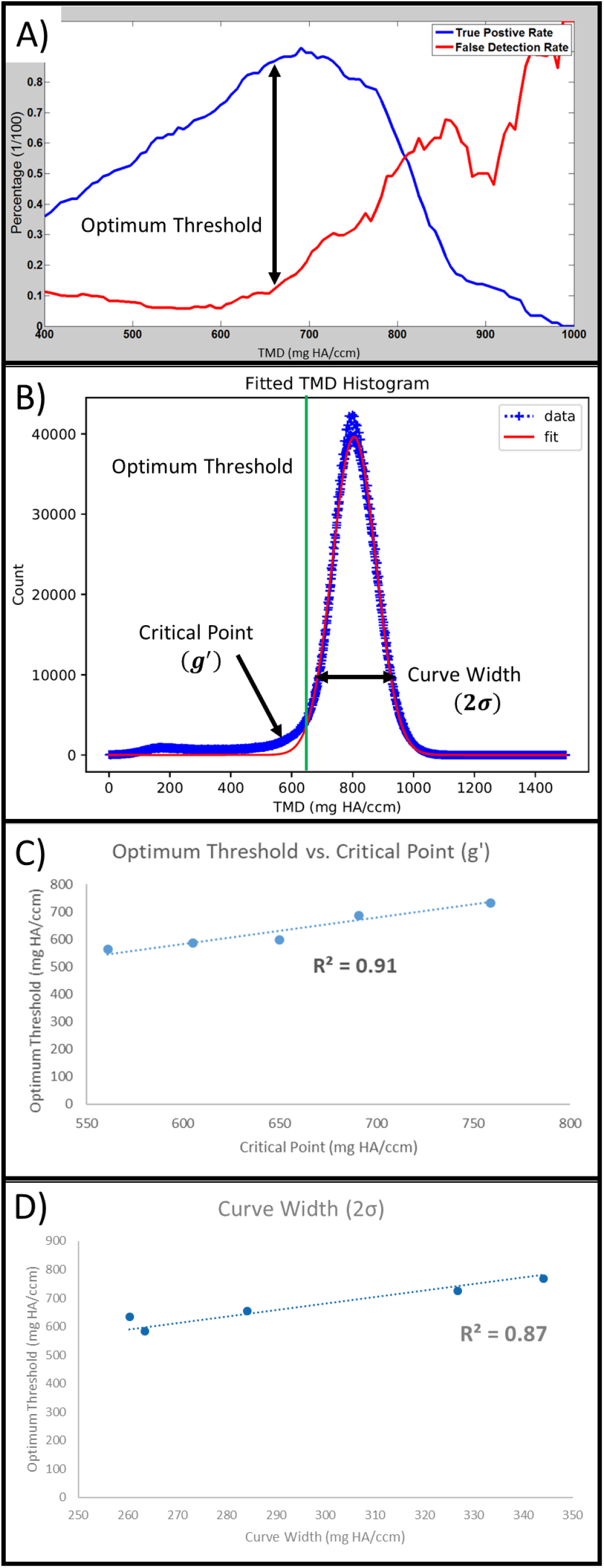
Individualized threshold selection. A) Iterative application of many single thresholds and each compared with the manual identification 3D coordinates of a given image subregion. Optimum threshold was defined as the single threshold that maximized the true positive rate and minimized the false detection rate. B) Typical TMD histogram of the bone biopsy’s micro-CT image. Optimum threshold (green) determined from (A) and the distribution characteristics including the critical point (g’) and curve width (2sigma) were calculated from the Gaussian fit of the data. C) Correlation between the optimum threshold for each of the five manually segmented subregions and the corresponding critical point (g’) from each respective TMD histogram. D) Correlation between the optimum threshold for each of the five manually segmented subregions and the corresponding distribution width (2sigma) from each respective TMD histogram.

**Figure S3:**
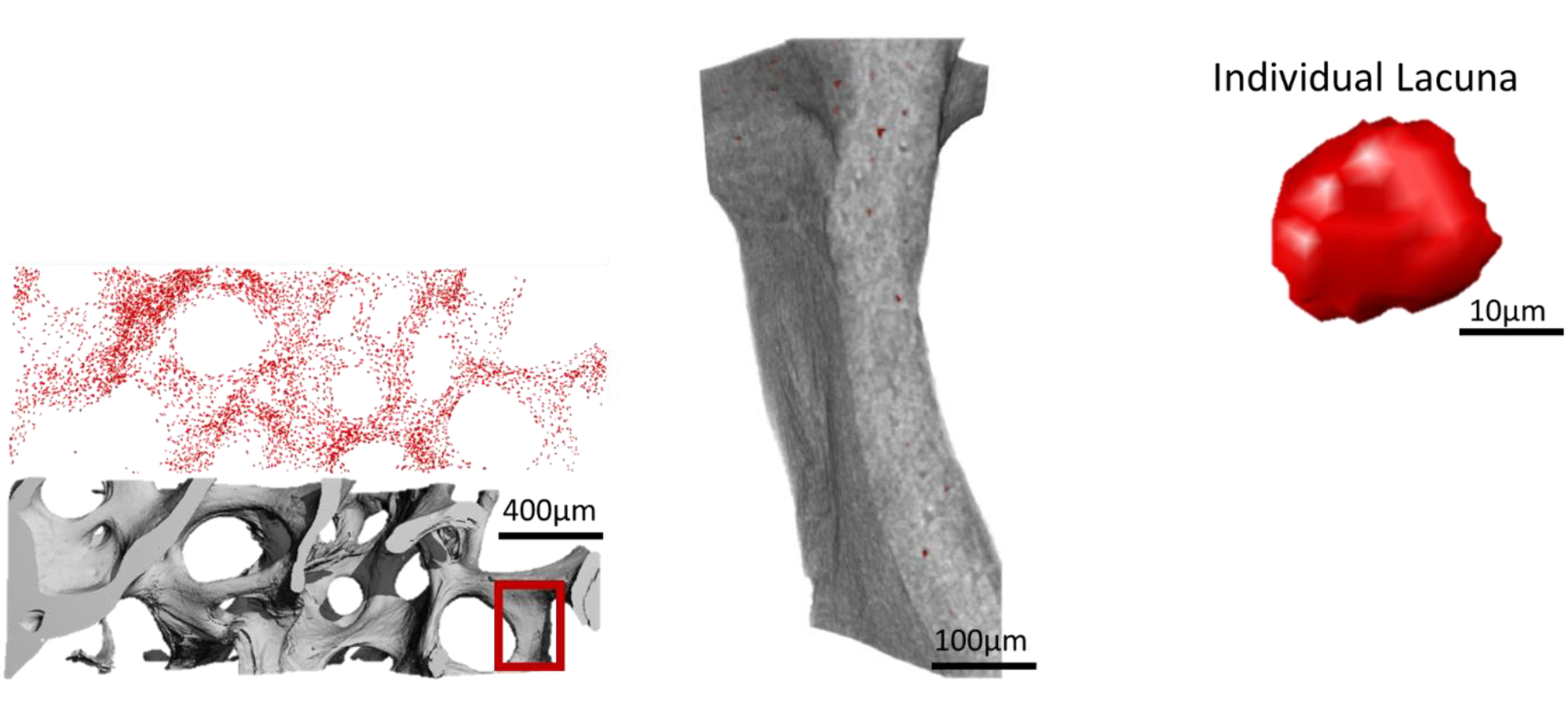
Animation of lacunar segmentation from the tissue level down to the individual cell level (see powerpoint file).

**Figure S4:**
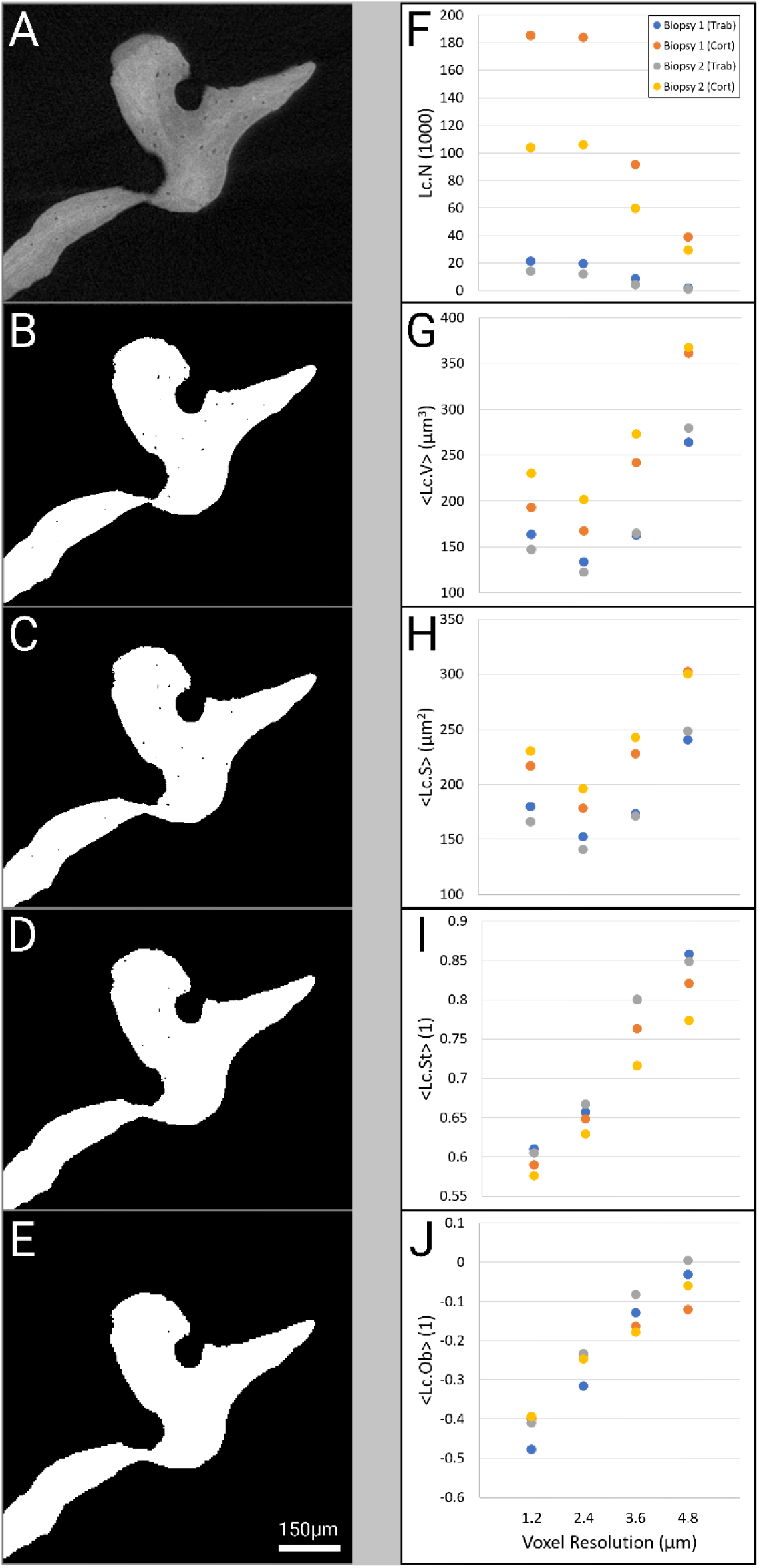
Lacunar morphometry resolution dependency. A) Original microCT grayscale filtered image of trabecular bone scanned at 1.2μm resolution. B) Thresholded image at original 1.2μm resolution. C) Thresholded image at downscaled 2.4μm resolution. D) Thresholded image at downscaled 3.6μm resolution. E) Thresholded image at downscaled 4.8μm resolution. Two region-matched biopsies (four regions in total) were analyzed at all four resolutions and morphometries plotted for F) lacunar number, G) lacunar volume, H) lacunar surface area, I) lacunar stretch, and J) lacunar oblateness.

