## Supplementary Material for "Large-scale quantification of human osteocyte lacunar morphological biomarkers as assessed by ultra-high-resolution desktop micro-computed tomography"

### Slide 1
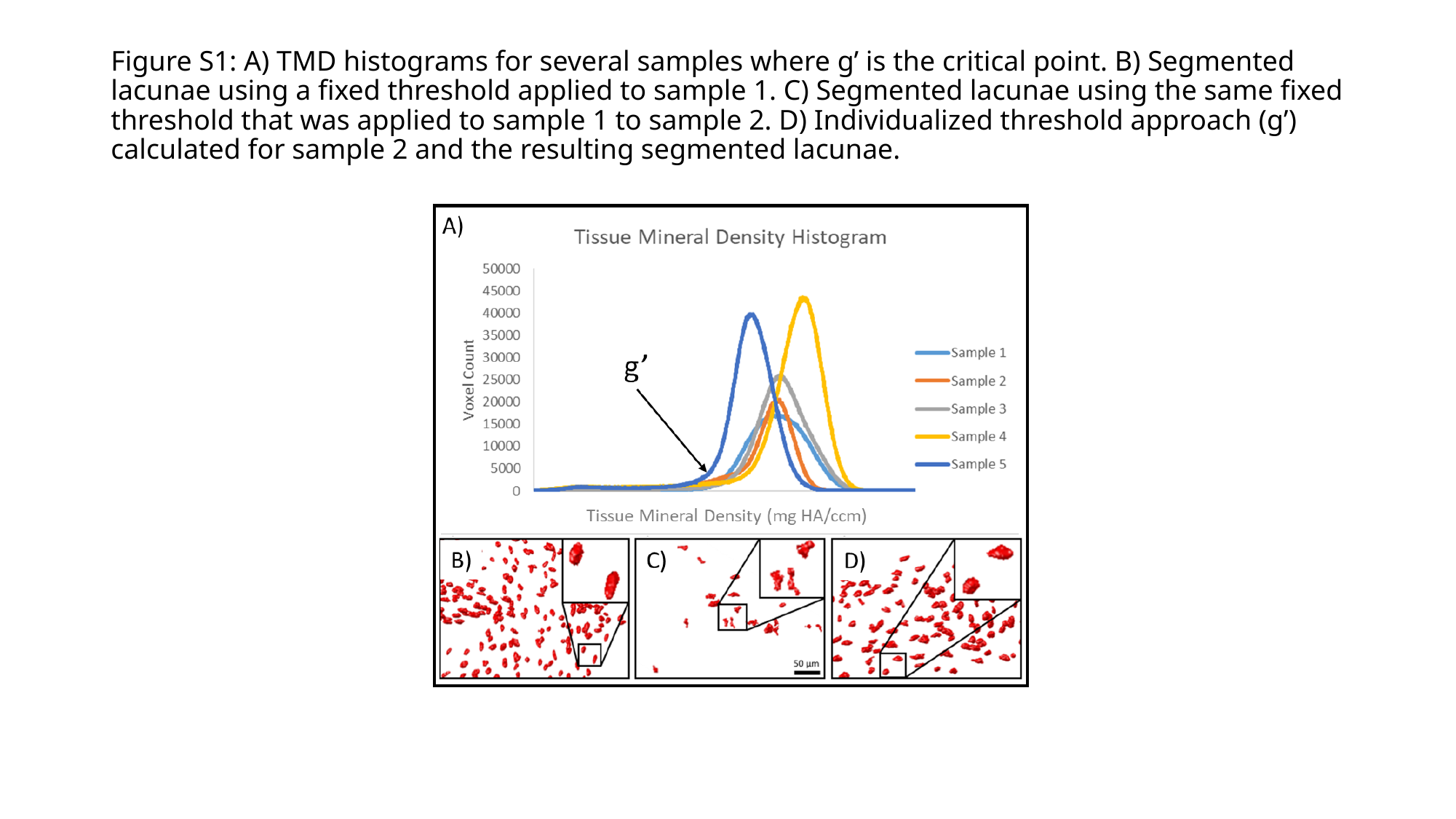

#### Figure S1: A) TMD histograms for several samples where g’ is the critical point. B) Segmented lacunae using a fixed threshold applied to sample 1. C) Segmented lacunae using the same fixed threshold that was applied to sample 1 to sample 2. D) Individualized threshold approach (g’) calculated for sample 2 and the resulting segmented lacunae.

### Slide 2
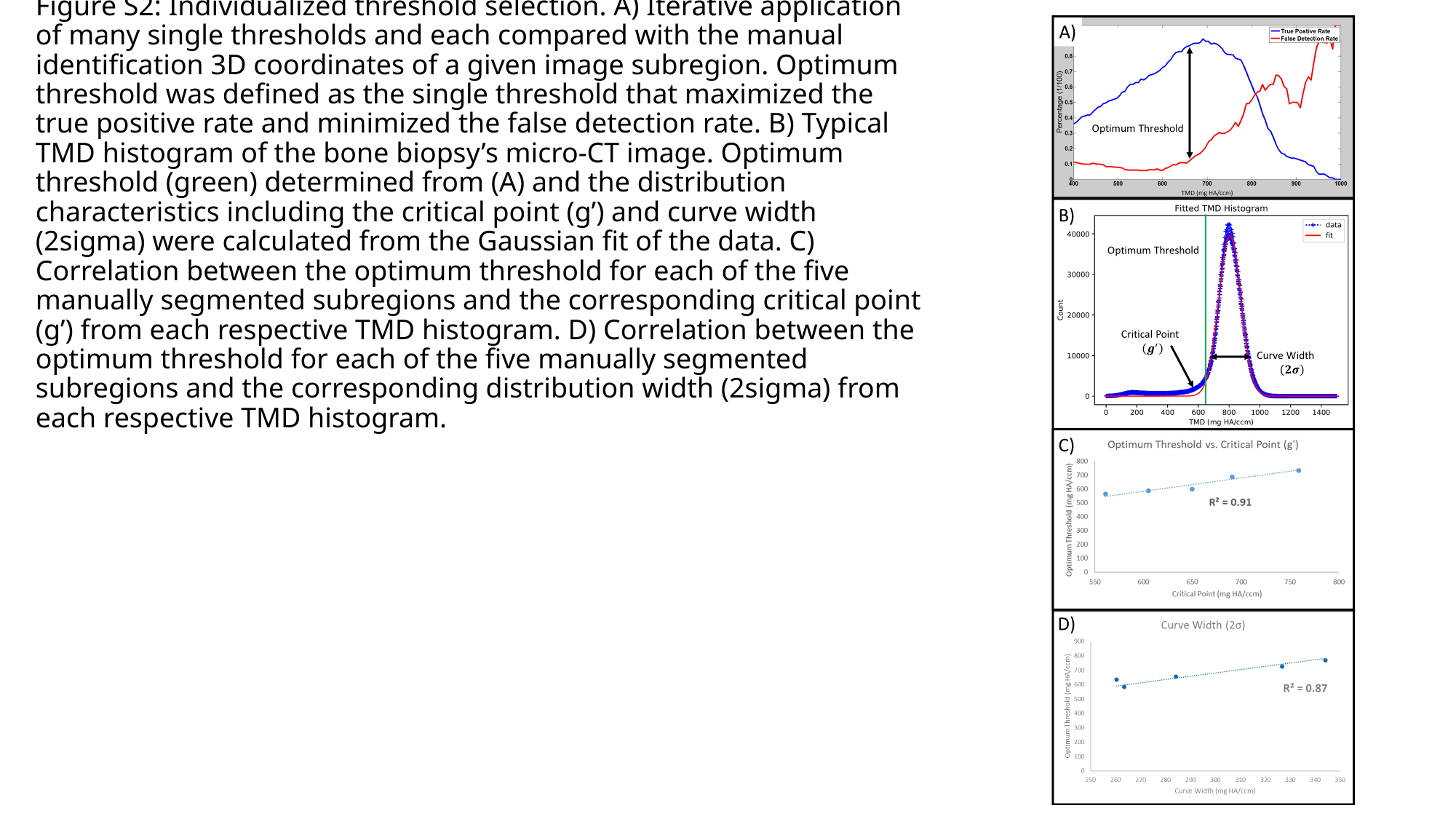

#### Figure S2: Individualized threshold selection. A) Iterative application of many single thresholds and each compared with the manual identification 3D coordinates of a given image subregion. Optimum threshold was defined as the single threshold that maximized the true positive rate and minimized the false detection rate. B) Typical TMD histogram of the bone biopsy’s micro-CT image. Optimum threshold (green) determined from (A) and the distribution characteristics including the critical point (g’) and curve width (2sigma) were calculated from the Gaussian fit of the data. C) Correlation between the optimum threshold for each of the five manually segmented subregions and the corresponding critical point (g’) from each respective TMD histogram. D) Correlation between the optimum threshold for each of the five manually segmented subregions and the corresponding distribution width (2sigma) from each respective TMD histogram.

### Slide 3
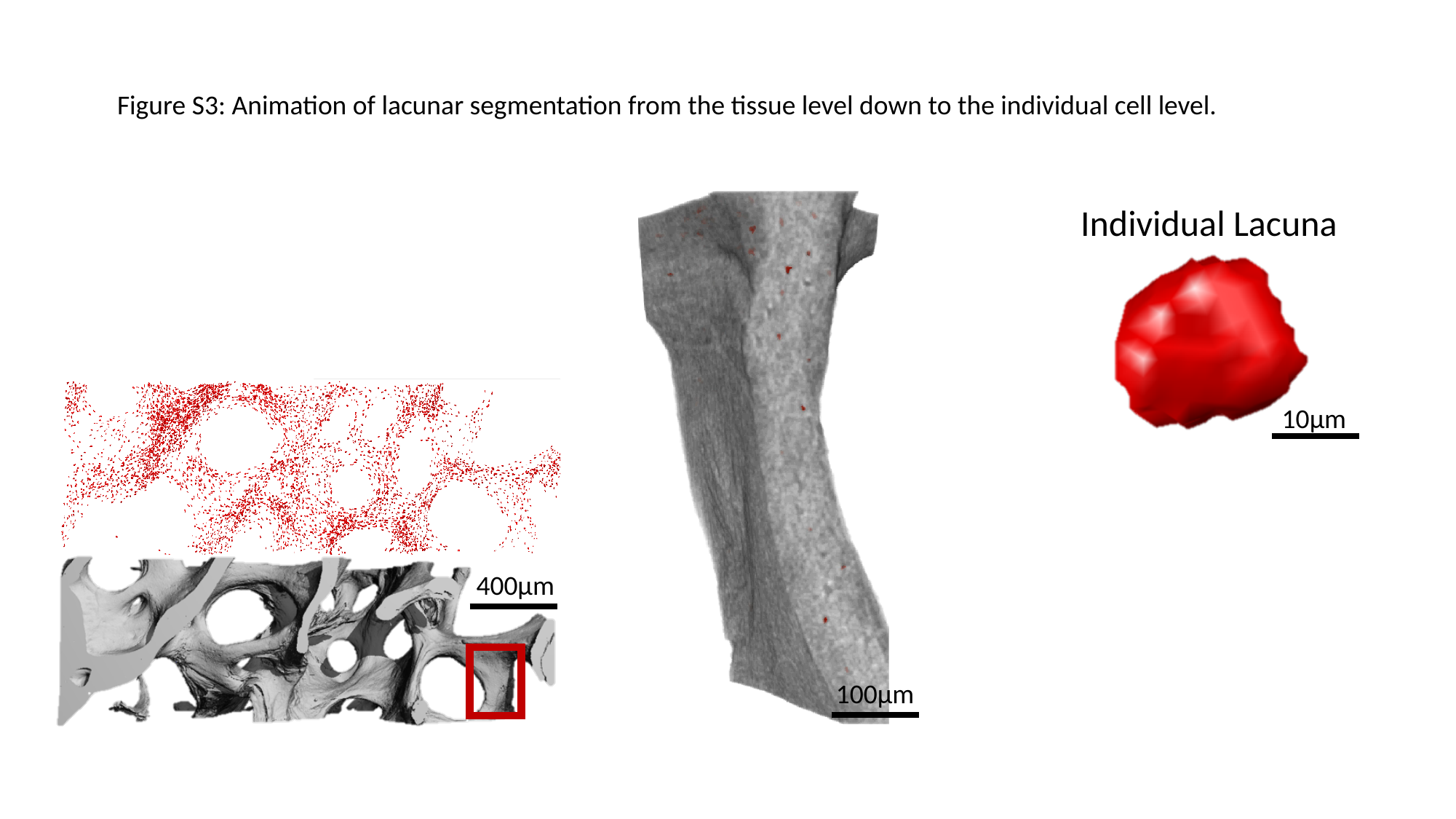

Figure S3: Animation of lacunar segmentation from the tissue level down to the individual cell level.
Individual Lacuna
10µm
400µm
100µm

### Slide 4
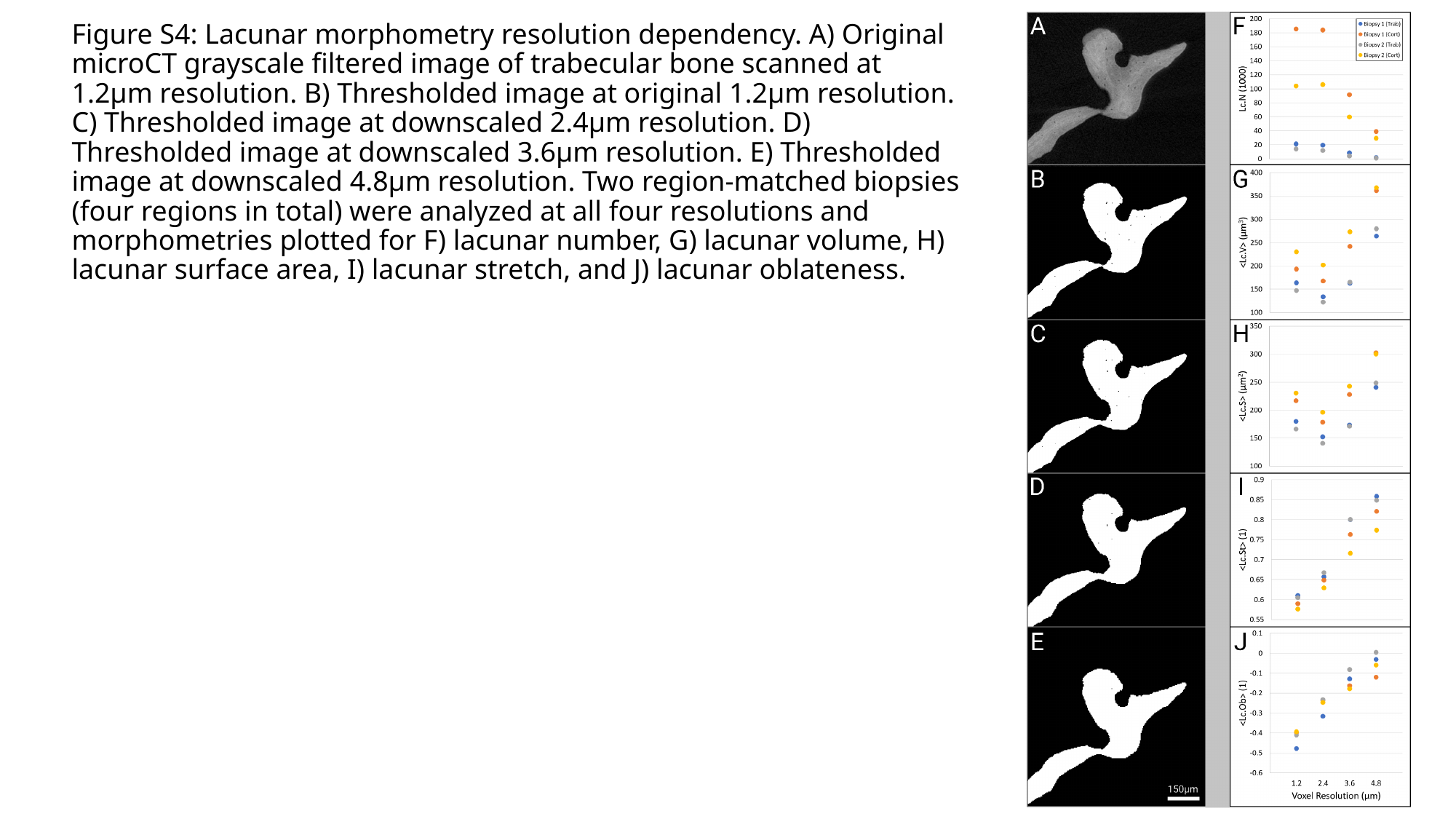

#### Figure S4: Lacunar morphometry resolution dependency. A) Original microCT grayscale filtered image of trabecular bone scanned at 1.2μm resolution. B) Thresholded image at original 1.2μm resolution. C) Thresholded image at downscaled 2.4μm resolution. D) Thresholded image at downscaled 3.6μm resolution. E) Thresholded image at downscaled 4.8μm resolution. Two region-matched biopsies (four regions in total) were analyzed at all four resolutions and morphometries plotted for F) lacunar number, G) lacunar volume, H) lacunar surface area, I) lacunar stretch, and J) lacunar oblateness.
